# 3D-Printed Autonomous Capillaric Circuits^†^

**DOI:** 10.1101/059238

**Authors:** A. O. Olanrewaju, A. Robillard, M. Dagher, D. Juncker

**Affiliations:** Biomedical Engineering Department, McGill University, 740 Dr Penfield Avenue, Montreal, QC, H3A 0G1, Canada.; McGill University and Genome Quebec Innovation Centre, 740 Dr Penfield Avenue, Montreal, QC, H3A 0G1, Canada; Department of Neurology and Neurosurgery, 740 Dr Penfield Avenue, Montreal, QC, H3A 0G1, Canada

## Abstract

Capillaric circuits (CCs) are advanced capillary microfluidic devices that move liquids in complex pre-programmed sequences without external pumps and valves-relying instead on microfluidic control elements powered by capillary forces. CCs were thought to require high-precision micro-scale features manufactured by photolithography in a cleanroom, which is slow and expensive. Here we present rapidly and inexpensively 3D-printed autonomous CCs. Molds for CCs were fabricated with a benchtop 3D-printer, Poly(dimethylsiloxane) replicas were made, and fluidic functionality was verified with aqueous solutions. We established design rules for 3D-printed CCs by a combination of modelling and experimentation. The functionality and reliability of 3D-printed trigger valves-an essential fluidic element that stops one liquid until flow is triggered by a second liquid-was tested for different geometries and different solutions. Trigger valves with geometries up to 80-fold larger than cleanroom-fabricated ones were found to function reliably. We designed 3D-printed retention burst valves that encode sequential liquid drainage and delivery using capillary pressure differences encoded by varying valve height and width. Using an electrical circuit analogue of the CC, we established circuit design rules for ensuring strictly sequential liquid delivery. We realized a 3D-printed CC with reservoir volumes 60 times larger than cleanroom-fabricated circuits and autonomously delivered eight liquids in a pre-determined sequence in < 7 min, exceeding the number of sequentially-encoded, self-regulated fluidic delivery events apreviously reported. Taken together, our results demonstrate that 3D-printing enables rapid prototyping of reliable CCs with improved functionality and potential applications in diagnostics, research and education.

## Introduction

Capillary-driven microfluidic devices move liquids using capillary forces defined by the geometry and surface chemistry of microchannels. This allows liquid delivery without using external pumps and valves. A wide range of capillary fluidic control elements were developed over the years including: stop valves,^1^ retention valves,^2^ trigger valves,^3^ and capillary pumps.^2,4,5^ Autonomous capillary microfluidic systems capable of self-powered and self-regulated completion of biochemical assays were also developed.^2,6–8^ Yet these autonomous capillary microfluidic systems were fabricated using silicon wafers and cleanroom processes with multiple photomasks, thereby increasing their cost and complexity. Paper-based microfluidics and lateral flow assays were also re-discovered as inexpensive approaches to autonomous capillary-driven flow;^9,10^ nevertheless, paper-based methods rely on heterogeneous porous substrates with statistical flow paths and cannot accomplish some of the valving capabilities that require the deterministic and predictable flow paths of microchannel-based devices. As such, there is a need for rapid and inexpensive fabrication of microchannel-based capillary microfluidics.

### Autonomous capillaric circuits for pre-programmed liquid delivery

More recently, advanced capillary microfluidic devices capable of pre-programmed delivery of multiple liquids were developed to enable autonomous multi-step processes, for instance to incorporate wash or signal amplification steps for improved bioassay sensitivity and specificity.^11–13^ Our research group proposed capillaric circuits (CCs) – advanced capillary circuits that are assembled from individual capillaric elements in the same way that electric circuits are assembled from individual electric components.^13^ CCs operate in a walk-away format where the operator pre-loads each reservoir, without worrying about the timing or sequence of these operations – instead, capillary microfluidic elements choreograph liquid delivery operations with minimal user intervention. This makes CCs a desirable platform for automating biochemical assays in point-of-care settings with minimal instrumentation.

Our group introduced two new fluidic elements to enable deterministic flow control with CCs. First, we developed two-level trigger valves (TVs) that stop liquids for over 30 minutes using an abrupt geometry change and a hydrophobic PDMS cover, thereby enabling pre-loading of reservoirs and subsequent liquid release when flow is triggered by a connected channel.^13^ We also developed retention burst valves (RBVs) that have a burst pressure encoded by their geometry. When integrated with other capillary fluidic elements within a CC, RBVs allow autonomous delivery of liquids in a pre-programmed sequence according to increasing order of RBV capillary pressure.^13^

### Rapid prototyping of capillary microfluidics

Although CCs enable sophisticated and automated fluidic operations, the prevailing view is that deterministic capillaric (and capillary) microfluidics require high-precision and small-scale (~10-μm) features for proper operation. As such, fabrication of CCs was dependent on cleanrooms, and was resource-intensive, time-consuming and expensive. Coupled with the need of photomasks for photolithography, a high cost and slow turnaround time for new design iterations limits the development of new devices and their widespread adoption.

To overcome the limitations of cleanroom fabrication, rapid and inexpensive prototyping of capillary microfluidic valves and integrated devices has been explored. An autonomous capillary system composed of capillary TV s, timing channels, and capillary pumps was fabricated by CO_2_ laser cutting.^14–16^ Although laser cutting enabled rapid prototyping (2-3 hours) of TVs that retain their liquid-stopping ability at widths up to 670 μm, and capillary pumps that retain their liquid-wicking ability with channel spacing up to 1120 μm, smaller features (≤ 300 μm) take on Gaussian geometries due to the laser manufacturing process. Refined elements, such as capillary retention valves, or more complex circuits have not been made using these techniques. Capillary stop valves have also been manufactured by injection molding, in a manner compatible with mass production (only 80 s to make each chip) but the passive valves were rendered hydrophobic to stop liquids and were not integrated within a self-powered and self-regulated capillary microfluidic system.^17^

### 3D-printed Microfluidics

Lately, there has been a surge of interest in 3D-printing for microfluidics applications due to the speed, accessibility, and low cost required to fabricate multilayer microfluidic structures. Recent reviews describe state of the art 3D-printing for microfluidics applications.^18,19^ All the demonstrations of 3D-printed microfluidics so far employ active flow control (usually pneumatic or centrifugal pumps). The resolution currently available with consumer grade 3D-printers is typically ≥ 200 μm^20,21^ with ~ 1 μm surface roughness.^22^ Capillary microfluidics however have traditionally been made with channels in the 1-100 μm range because the capillary pressure is inversely proportional to the smallest dimension, and becomes very small for large microchannels. Moreover, valving and flow control depend on the surface topography and abrupt geometric changes and low surface roughness are considered necessary to prevent pre-wetting and creeping flows. Hence the prevailing perception is that current 3D-printing technology may not be suitable for making capillary microfluidics because the smallest dimensions are too large to obtain adequate capillary pressure, the resolution and precision insufficient for making abrupt changes needed for reliable valves – notably due to the layered structure of stereolithographic printing forming steps that lend themselves to corner flow-and the high surface roughness may lead to creeping of liquid.

### 3D-printed Capillaric Circuits

Here we present complex capillaric microfluidic circuits made by stereolithographic 3D-printing with geometries scaled up > 20-fold. 3D-printing allows rapid and inexpensive fabrication of CCs. This enables investigation and engineering of CCs with greater capabilities and increased accessibility in research and point-of-care settings. First, we 3D-print TVs and characterize their performance as a function of geometry and surfactant concentration. Then we investigate design rules for 3D-printed CCs composed of TVs, RBVs, flow resistors, and capillary pumps using a proof of principle circuit with four reservoirs. Finally, we demonstrate the capabilities of our 3D-printed capillarics by developing a circuit for autonomous delivery of eight liquids in < 7 minutes.

## Materials and Methods

### Process flow for 3D-printing capillaric circuits

First, we developed a symbolic representation for CCs using electrical analogies, as described in our previous work.^13^ Next, the symbolic circuit was converted into a computer-aided schematic design that was exported into the standard stereolithography (STL) format for 3D-printing. We 3D-printed molds (negatives) of capillaric microfluidic devices using a stereolithography-based printer (Perfactory MicroEDU, EnvisionTEC Inc., USA) with 96 μm XY pixel size and 50 μm Z layer height. Microfluidic features were aligned to the pixel grid of the 3D-printer projector to ensure accurate realization of features. The 3D-printer’s default settings were used. Device designs included 2 mm thick bases for easier handling. The typical printing time for capillaric microfluidic devices, with multiple devices arranged to cover nearly the entire 100 × 75 mm^2^ print area of the 3D-printer, was ~30 minutes. After 3D-printing, molds were washed in isopropanol for 5 minutes and dried with nitrogen gas. 3D-printed molds were inspected under the microscope to check for defects during the printing process.

### PDMS replication from 3D-printed molds

To obtain multiple copies of capillary microfluidic devices from the same 3D-printed mold, we made Poly(dimethylsiloxane) (PDMS) replicas of devices by soft lithography.^23^ We 3D-printed molds using a high temperature molding resin (HTM140 resin, EnvisionTEC Inc., USA) with a manufacturer-specified heat deflection temperature of 140°C to allow replica molding of 3D-printed structures. Prior to PDMS replication, molds were pre-treated with a silicone spray (Ease Release 200®, Mann Formulated Products, USA) to prevent PDMS from sticking to the mold. The spray was applied in two passes uniformly over the surface of the mold from a height of about 10 cm. To make PDMS replicas, elastomer base and curing agent (Sylgard 184, Paisley Products Inc., Canada) were mixed in a 10:1 ratio. The PDMS mixture was degassed for 1 hour and poured onto the 3D-printed mold placed in a petri dish. PDMS was cured overnight at 60°C and then peeled from the mold. First PDMS replicas were discarded because they were sticky due to the presence of silicone spray residue; subsequent replicas were used for capillary microfluidics experiments.

### Procedure for capillary-driven flow experiments

To obtain hydrophilic surfaces for capillary-driven flow, PDMS replicas were activated for 12 seconds at 200 mTorr & 150 W in a plasma chamber (PE-50, PlasmaEtch, USA). To characterize the plasma-treated surfaces, advancing and receding contact angles of deionized water were measured using a video-based optical contact angle measurement instrument (OCA 15EC, Dataphysics Instruments GmbH, Germany). Plasma-treated PDMS devices were sealed with flat, untreated PDMS covers to provide closed microchannels for capillary-driven flow. The PDMS covers were made with a 1:20 ratio of curing agent to elastomer base to obtain soft and flexible PDMS surfaces that sealed well, despite their hydrophobicity. Flow in CCs was tested using aqueous food dye solutions and visualized under a stereomicroscope (SMZ-8, Leica Microsystems Inc., Canada) with a video camera (Lumix GH3 DSLR, Panasonic Inc., Canada). During TV testing, when devices were tested for ≥ 30 min, we humidified the area around the capillary microfluidic chips with wet Kimwipes^®^ and covered with a petri dish to prevent evaporation.^24^

## Results and Discussion

CCs operate using a series of functional elements including inlets, channels, flow resistors, capillary pumps, trigger valves (TV), capillary retention valves, and retention burst valves (RBVs) that can be combined for encoding the pre-programmed delivery of multiple liquids.^13^. The capillary pressure of each RBV is calculated using the Young-Laplace equation:

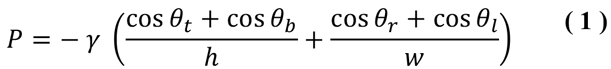

where *P* is the capillary pressure, *γ* is the surface tension of liquid in the microchannel, and *h*, *w*, are the channel height and width respectively. *θ_t_*, *θ_b_*, *θ_r_*, *θ_l_*, are the top, bottom, right, and left channel wall contact angles, respectively. Contact angle hysteresis must be taken into account when designing RBVs since the advancing contact angles are relevant when a channel is filled while the receding contact angles are relevant when a channel is drained. Likewise, the resistance *R* for a conduit with a rectangular cross-section is given by:^25^

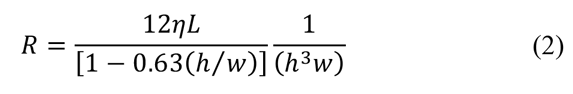

Where *η* is the viscosity of liquid in the channel, and *L* is the length of the microchannel. The cross section of channels and various elements for microfabricated CCs reported by Safavieh *et al*^13^ ranged from 15 × 100 μm^2^ to 200 × 200 μm^2^. Thus, assuming receding contact angles of 89° and 31° for the hydrophobic top PDMS surface and hydrophilic side and bottom surfaces respectively, and a surface tension *γ* of 72 N/m, the capillary pressures of microchannels (calculated using equation 1) in the cleanroom-fabricated circuits ranged from – 7948 Pa to – 1264 Pa. These capillary pressures would correspond to water column heights of 810 mm and 129 mm respectively in capillary rise experiments. Since our microchannel lengths were on the order of 5 mm, capillary forces dominated gravity in our microfabricated CCs and our devices could be operated without considering gravity effects.

We first tested whether 3D-printed channels and capillary pumps replicated into PDMS could be filled by a liquid, and found that this worked reliably up to 1000 × 1000 μm^2^ constituting the upper limit for capillary elements in this study. The lower size limit for fluidic elements was set by the resolution of the 3D printer. The vertical resolution was set by the thickness of each printed layer and was 50 μm. The lateral resolution was 100 μm under the best circumstances, but was limited to 200 μm when taking into consideration fabrication yield. Hence, for 3D-printed circuits, the cross-sectional dimensions range from 50 × 200 μm^2^ to 1000 × 1000 μm^2^, and the capillary pressure ranges from – 955 Pa to – 188 Pa, or a water column height from 97 mm to 19 mm. These type of conduits filled spontaneously with aqueous solutions and remain in a microfluidic regime where gravity and inertia within the conduits are negligible.

Next, we set out to test whether critical functional elements such as the TV and the RBV could also be 3D printed, whether surface roughness might affect their functionality, and to determine the design rules for making them.

### Trigger Valves

In order to develop functional CCs, the first step is to have functional and reliable trigger valves (TVs) to robustly hold liquids in reservoirs.^13^ Consequently, we first characterized 3D-printed TVs on a standalone basis, before developing more complex CCs.

#### Cleanroom-fabricated versus 3D-printed trigger valves

Cleanroom fabrication is generally considered the gold standard for manufacturing capillary stop valves and TVs because of the small feature sizes and smooth channel surfaces attainable.^2,5,6,11–13,26,27^ Cleanroom-fabricated TVs have small features (~ 20 μm) and smooth, vertical channel walls (Fig. 1a & 1c). Meanwhile, 3D-printed trigger features have larger minimum widths (≥ 100 μm) and rough, layered channel walls (Fig. 1b & 1d). These stark geometry differences call into question the functionality and reliability of 3D-printed TVs.

**Figure 1.**
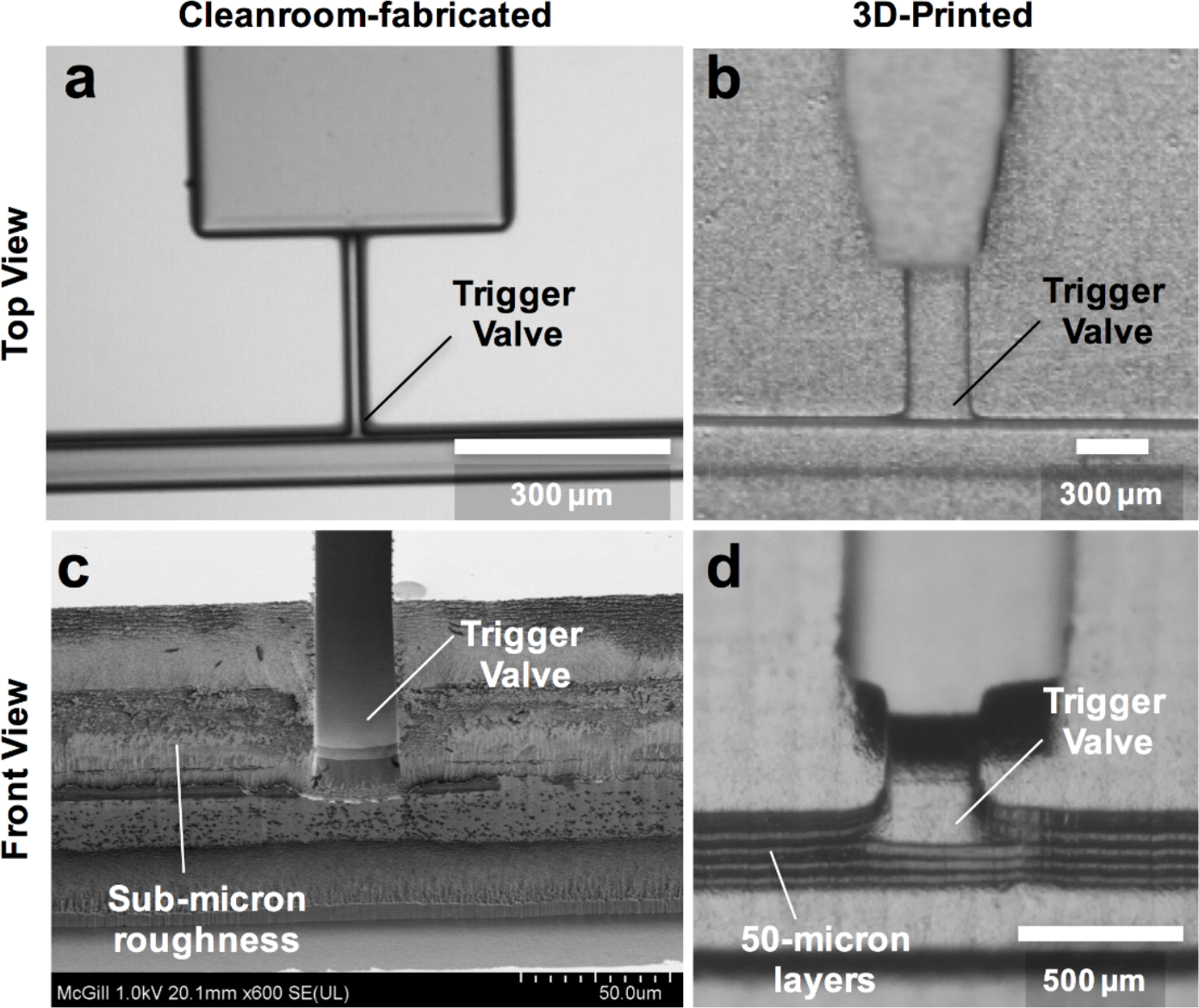
Comparison between cleanroom-fabricated and 3D-printed TVs. a) Top view of TV fabricated by photolithography in the cleanroom showing smooth, high-precision features. b) Top view of TV fabricated by stereolithography-based 3D printing showing rough, large features. c) Scanning electron micrograph of TV fabricated by deep reactive ion etching of silicon showing vertical channel walls with sub-micron roughness. d) Front view of PDMS replica of 3D-printed TV showing 50-μm thick ridges on the channel wall due to the layer-by-layer printing process.

We 3D-printed TVs and tested a wide range of geometries and surfactant concentrations to assess their functionality and reliability. Previously, the success rate of capillary stop valves and TVs was only reported over a 5-minute period.^16,26^ Here we defined TV success as when a valve holds liquid for at least 30 minutes without leakage. This allowed autonomous microfluidic operations in a walk-away format where the user pre-loads samples and reagents onto the chip and subsequently starts the assay at a time of their choosing, without needing to fit their operations to a strict 5-minute window.

#### Effect of trigger valve geometry on success rate

The geometry of TVs influences their success rate.^17,26,28^ Fig. 2a shows the geometric parameters known to affect the performance of capillary TVs: the height of the TV, width of the TV, and the height difference between the TV and its release channel. To determine which geometries provide high TV success rates, we tested valves with widths of 96 μm, 192 μm, 288 μm, 480 μm, 672 μm, 960 μm, and 2016 μm. TV heights were fixed at either 400 μm or 1000 μm to obtain different height-to-width ratios for these experiments. As summarized in Fig. 2b, all TVs tested were at least 75 % successful (N=8). The few failures were due to difficulties while loading valves with low (< 1) or high (> 5) aspect ratios (i.e. height-to-width ratios) that required the user to apply additional positive pressure when filling the valves. We found that 3D-printed TVs were reliable with dimensions up to 3 times larger than reported with CO_2_ laser cutting^14^ and up to 80 times larger than typical cleanroom-fabricated valves.^13,26^

**Figure 2.**
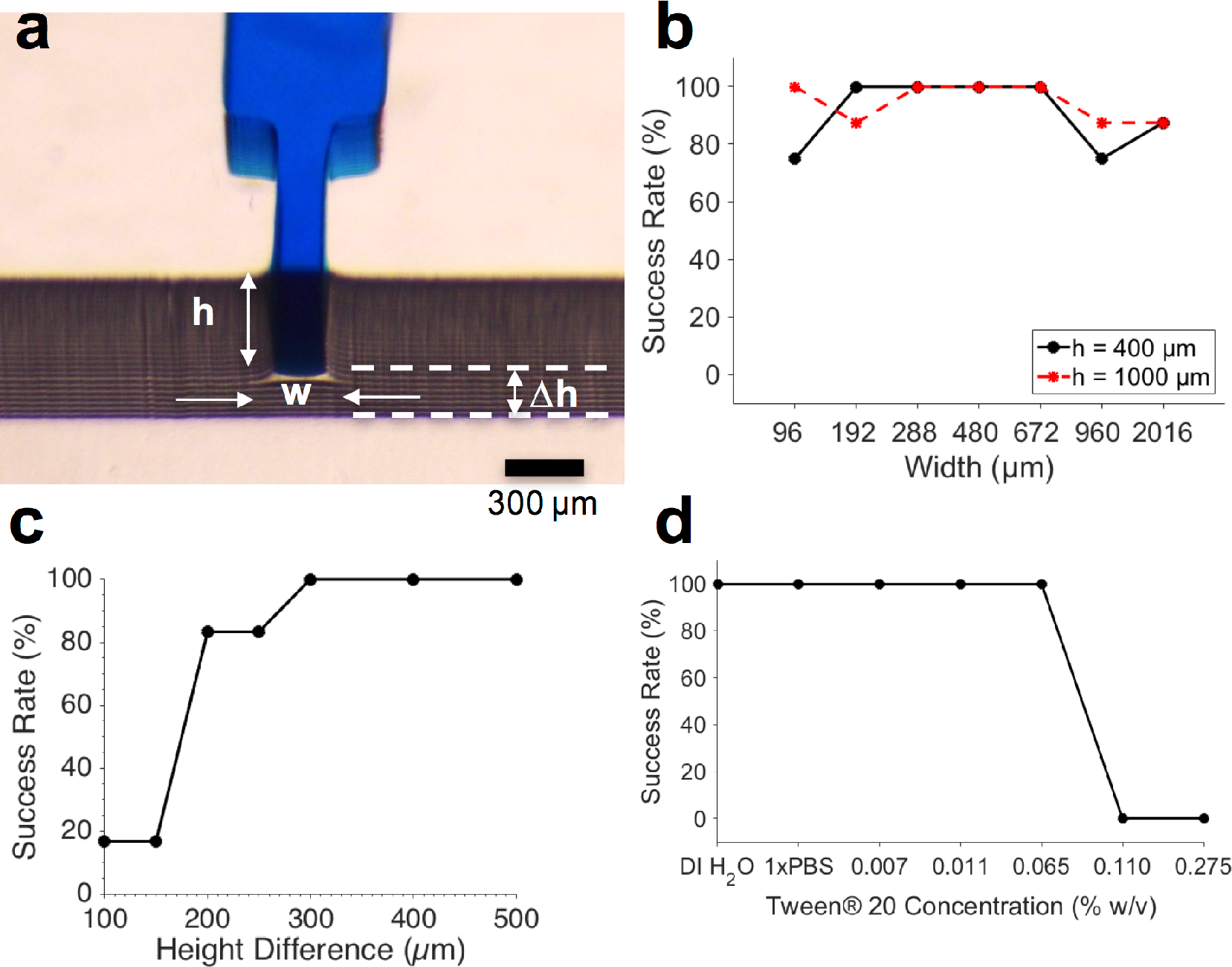
Effect of geometry and surfactant concentration on success rate of 3D-printed TVs. a) Front view of food dye solution stopped at 3D-printed TV showing the TV height (h), width (w), and the height difference between TV and release channel (Δh). b) Success rates for 3D-Printed TVs over a wide range of widths and heights. N = 8. c) Above a height difference (Δh) of 300 μm, 3D-Printed TVs were 100 % successful. N = 6. d) TVs were 100 % successful at Tween^®^ 20 concentrations ≤ 0.0650 % weight/volume in 1xPBS. N = 3

Since the minimum z-layer thickness of the microchannels was limited to 50 μm by the 3D-printer resolution, we tested height differences of 100 μm, 150 μm, 200 μm, 250 μm, 300 μm, 400 μm, and 500 μm between the TV and the release channel. The TVs used for these height difference tests were 300 μm wide and 50 μm deep, since our TV characterizations showed reliable functionality over a wide range of geometries (Fig. 2b). As seen in Fig. 2c, the height difference between the TV and the release channel had a threshold effect on TV success. When the height difference was ≥ 300 μm, TVs were 100 % successful (N = 6).

#### Effect of surfactant concentration on trigger valve performance

Despite the fact that the most common application of capillary microfluidics is to automate biological assays that often require the use of surfactant-containing reagents, the effect of surfactant concentration on TV performance is not well reported in the literature. To determine the effect of surfactant on TVs, we tested aqueous solutions with different concentrations of Tween^®^ 20, a surfactant commonly used in immunoassay wash buffers and for cell lysis. The critical micelle concentration of Tween® 20, is 0.0074% w/v. Consequently, we tested the following concentrations of Tween^®^ 20: 0.0074, 0.0110, 0.0650, 0.1100, and 0.2750 % weight/volume. As shown in Fig. 2d, we found that the TVs were 100 % reliable when Tween^®^ 20 concentrations were ≤ 0.0650 % weight/volume (N=3), a suitable surfactant concentration for use in wash buffers during immunoassays that commonly use 0.05%.^29,30^

### Retention Burst Valves

Retention burst valves (RBVs) retain liquid in a conduit up to a threshold, or bursting, pressure which if exceeded leads to bursting of the valve and draining of the liquid held downstream in a reservoir. It is thus possible to drain a series of reservoirs connected to a main channel in a predetermined sequence by terminating each of them with a RBV with increasing burst pressure. The burst pressure of a RBV can be calculated using Eq. 1 and using the receding contact angles for the liquid which were found to be 95° for the hydrophobic PDMS cover, and 31° for the hydrophilic bottom and side walls.

#### Capillaric circuit for autonomous delivery of four liquids

As a proof of principle of 3D-printed capillarics, we designed a circuit with 4 RBVs (Fig. 3a & b). PDMS replicas of the 3D-printed mold were made (Fig. 3c), plasma-treated for hydrophilicity, and sealed with a hydrophobic PDMS cover (Fig. 3d). The expected pre-programmed operation of the CC is illustrated in Fig. 3e. First reservoirs were filled and TVs held each liquid in place. Next, a solution was added to the release channel, connecting the reservoirs to the pump and starting the preprogrammed liquid delivery sequence. Subsequently, the RBVs burst sequentially according to increasing capillary pressure.

The TVs in the CC were designed to have the smallest cross section in the circuit and the highest capillary pressure in the CC since they play a dual role – stopping liquids during initial filling of reservoirs, and acting as retention valves with higher capillary pressure than the capillary pump during reservoir drainage (see Figure 3a). These retention valves ensure that the side branches are not completely emptied (with minimal dead volume), thereby allowing sequential liquid delivery without bubble trapping. ^2,13^

In cleanroom-fabricated devices, multiple masks are needed for making structures with multiple depths; hence only the microchannel widths were used as a free parameter to adjust the RBV threshold.^13^ RBVs were typically 100 μm deep and had widths of 200 μm, 130 μm, 110 μm, and 90 μm corresponding to capillary pressures of – 1264 Pa,–1601 Pa,–1847 Pa, and–2028 Pa respectively.

With 3D-printing both the width and depth can be adjusted independently and fabricated in one shot. Consequently, we encoded the capillary pressure differences between 3D-printed RBVs by modifying both the height and widths of the microchannels. The lower size limit of our microchannels was set by 3D-printer resolution. The pixel size for the EnvisionTEC Perfactory MicroEDU 3D-printer is listed as 96 μm, the smallest features that we were able to print with a high yield were 200 μm wide and 50 μm deep open channels. The resolution obtained is similar to that reported for other state of the art stereolithographic 3D-printers in the literature.^18^

**Figure 3.**
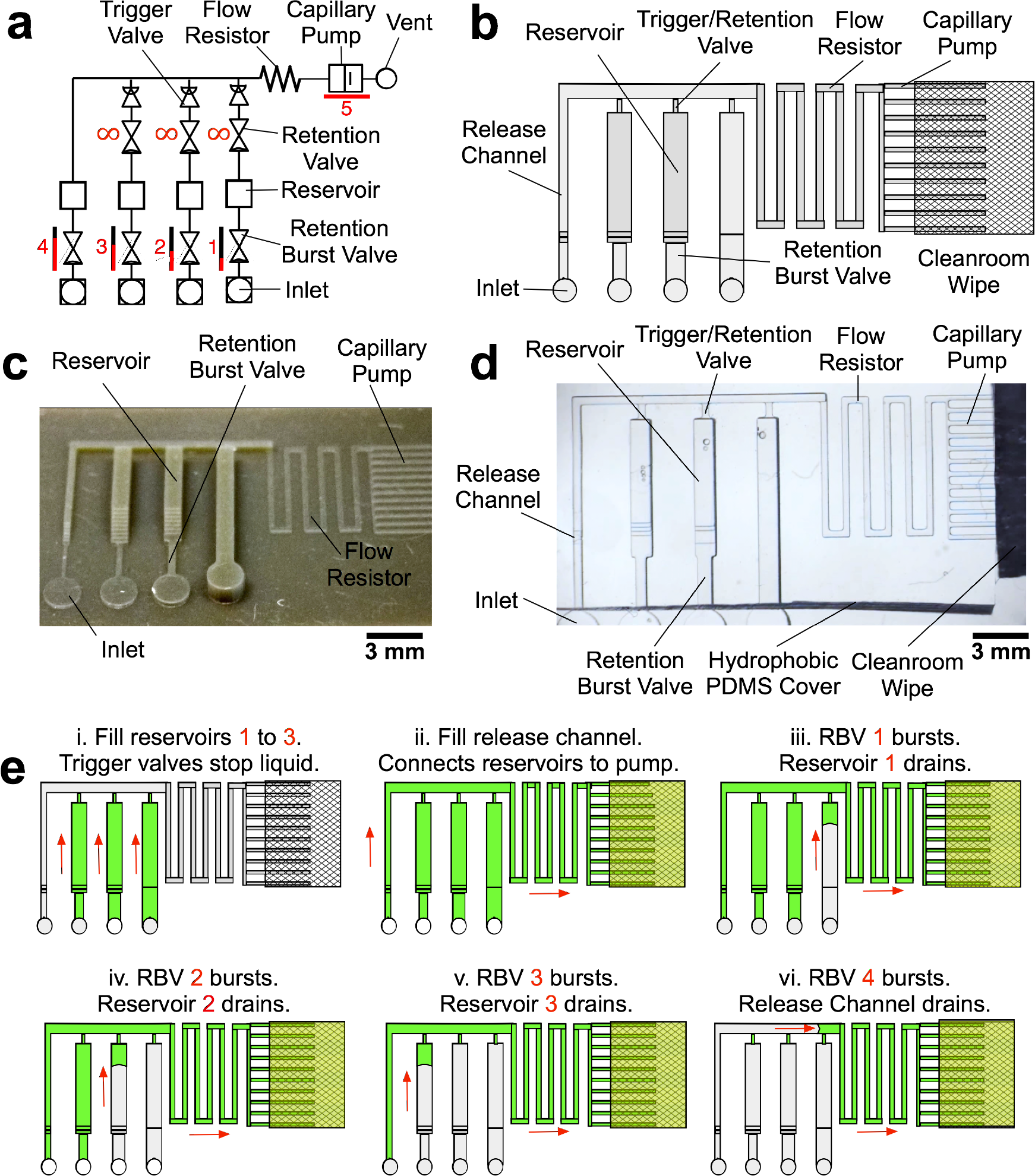
Design, mold, PDMS replica and operation of 3D-printed CCs for autonomous sequential delivery of four liquids. a) Symbolic representation of CC with main fluidic elements labelled. b) Schematic of CC. c) 3D-printed mold of the CC and d) PDMS replica with transparent PDMS cover and clean room wipe contacting the capillary pump. e) Schematic illustrating expected operation of RBVs. Solutions loaded into the reservoirs are delivered in pre-programmed manner according to RBV capillary pressure.

The 3D-printed reservoirs were 960 μm wide, 1,000 μm deep, and 6250 μm long with a volume of 6 μL, corresponding to 60 times the volume of typical microfabricated reservoirs.^13^ Due to the change in the size of the RBV, there were minor changes in volume for each reservoir (Fig. 3c & d) that could be compensated for by adjusting the reservoir size.

To accommodate the large volumes in the reagent reservoirs without significantly increasing device footprint, we placed a cleanroom wipe made of paper (Durx 670, Berkshire Corporation, USA) atop the 3D-printed capillary pump.^31^ The combination of 3D-printing and off-the-shelf, low cost paper saves costs without compromising performance. The driving capillary pressure in the circuit is defined by capillary pump because the gap between the edge of the PDMS and the paper forms an open microchannel that can be drained. Hence, the capillary pressure is dictated by the capillary pressure of the capillary pump, allowing use of paper pumps with higher, but sometimes ill-defined capillary pressures, without impacting the accuracy and functionality of the CCs.

**Figure 4.**
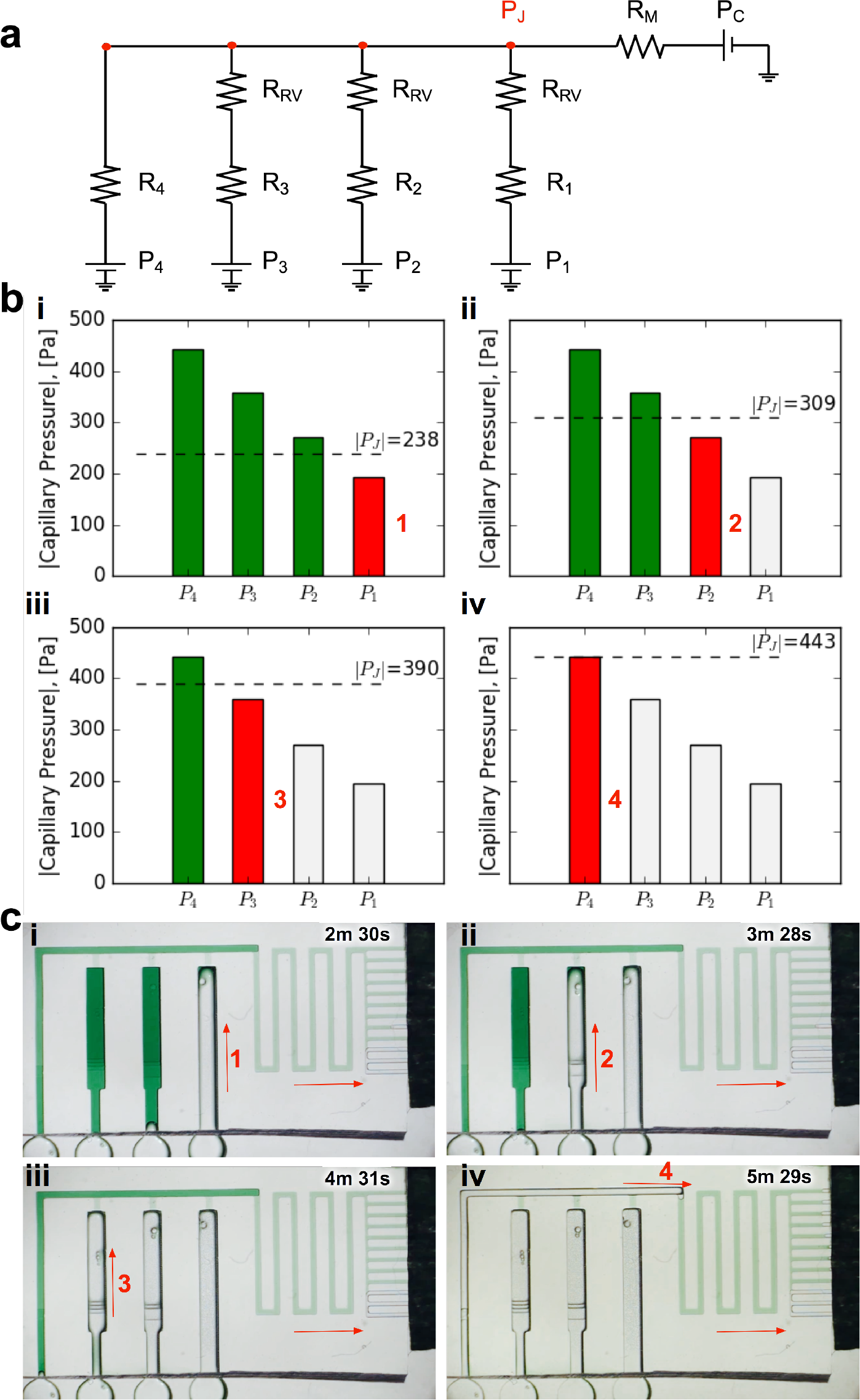
Design and experimental validation of CC for autonomous delivery of four liquids. a) Electric circuit analogue showing the flow resistances and capillary pressures in the CC. b) Graphs showing the calculated junction pressures during bursting of each RBV. Junction pressures were designed to ensure that RBVs burst sequentially. c) Time-lapse images showing autonomous and sequential drainage of reservoirs in the CC. Arrows represent sequence and flow direction. Text labels show time during liquid delivery. A video of the autonomous liquid delivery operation is provided in supplementary movie S1.

#### Requirements for sequential RBV bursting

It is not sufficient to simply increase the burst pressure of the RBV to achieve sequential drainage and in fact the architecture of the CC must be designed to ensure that an RBV only bursts after complete drainage of the reservoir connected to the previous RBV. To illustrate this point, the proof of concept CC shown in Fig. 3a when filled with liquid is modeled by an electrical equivalent circuit shown in Fig. 4a.

Considering the circuit at the instant when all reservoirs are filled, but still under static conditions, without flow, the junction pressure PJ will be equal to the pressure PC of the capillary pump. Given that the capillary pressure of the pump is larger than the capillary pressure of the side branches, liquid will be drawn towards the junction PJ, leading to flow in the CC. The first side branch to be drained in the CC is the one connected to the RBV with the lowest burst pressure, which here is RBV1 with pressure Pi. As liquid drains from side branch 1, there is a pressure drop across Ri and RRV on one hand and across the main resistor RM on the other hand, which will lead to a reduction of the pressure PJ at the juncture between the side-branch and the main channel. The high resistance of RRV and RM compared to the low resistance of the release channel ensures that pressure PJ is replicated across all 4 junctions (red dot in Fig. 4a). To avoid bursting of RBV2 while branch 1 is draining, it is imperative that |P_J_| < |P_2_| at all times. Assuming a single branch drains at any given time, the pressure PJ during drainage of the RBV can be calculated from the electrical circuit analogue using Kirchhoff’s law and Ohm’s law yielding:

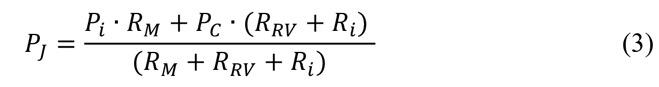

where the index *i* represents the side branch that is being drained in the capillary circuit, *P_i_*is the capillary pressure of the liquid meniscus on the end of the side branch, *R_i_*is the flow resistance of the RBV and reservoir of the side branch, *R_RV_* is the resistance of the retention valve, *R_M_* is the flow resistance of the main resistor, and *P_c_* is the pressure of the capillary pump (see Fig. 4a).

As the liquid drains, the resistance in the side branch is expected to change. However, the retention/trigger valve structure has the smallest cross-section (300 × 50 μm^2^) in the side branch and its associated resistance R_RV_ > 100 R_i_. Consequently, changes in the resistance of the side branch during drainage are negligible and do not need to be considered in the calculation of P_J_. Moreover, after the RBV drains, P_J_ decreases since the capillary pressure at the end of the side branch now becomes the capillary pressure of the reservoir rather than the capillary pressure of the RBV. This drop in P_J_ does not adversely affect sequential liquid delivery since the condition for sequential liquid delivery is still met; in fact, during reservoir drainage one expects liquid delivery to be even more sequential since the junction pressure is lower. Hence the most stringent condition on PJ is given by the situation described by the electrical circuit with fully filled conduits, which can thus be used to establish the conditions for sequential drainage of each of the side branches.

The required condition for junction pressure to ensure that only one RBV in the CC bursts can be generalized as follows:

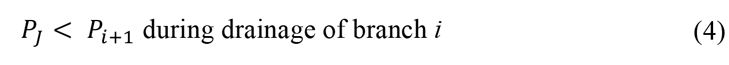

where *P*_*i*+1_ is the capillary pressure of the next RBV to burst in the circuit. This condition can be satisfied by balancing the flow resistance in the circuit, and in particular adjusting the main flow resistance (R_M_) in front of the capillary pump to ensure that P_J_ during drainage is lower than the pressure of the subsequent RBVs (see Fig. 4a). This calculation is applicable to CCs where the resistance of the channel linking the side branches is negligible compared to R_RV_, or else that resistance must also be considered and the appropriate analogous electrical model derived and resolved. The calculation holds for the model CC and can be used to calculate the pressure P_J_ during drainage of branch i and ensure that it is smaller than the retention pressure P_i+1_ of branch i+1 Fig. 4b.

The geometries of the RBVs in the CC are summarized in Table 1. We designed our proof-of-principle device to obtain uniform capillary pressure differences of 80 ± 5 Pa between successive valves. All RBVs were 2.6 mm long. We designed a 4.2 mm long, 290 μm wide, and 100 μm deep main resistor so that the junction pressure during each liquid delivery step satisfied our condition for sequential liquid delivery (see Fig. 4b).

Contact angle hysteresis must be taken into account when designing the capillary pump to ensure that the capillary pressure threshold for all RBVs can be overcome. Since the filling of the capillary pump is dictated by the advancing contact angles on the microchannel walls while the bursting of the RBVs is dictated by the receding contact angles on the microchannel walls, the dimensions of the capillary pump must be significantly smaller than the smallest dimension of RBVs to ensure drainage.^13^ Thus, the capillary pump in our proof-of-principle 4-valve circuit was using microchannels that were 200 × 100 μm^2^, providing a wicking capillary pressure of-736 Pa that is large enough to drain each RBV in the circuit.

**Table 1.**
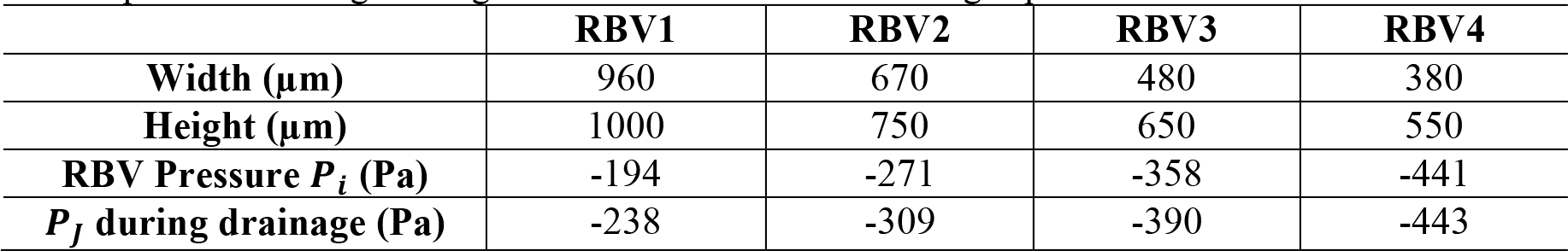
Geometry of retention burst valves (RBVs) in CC for autonomous delivery of four liquids. Junction pressures during drainage of RBVs were calculated using Equation 3.

|  | **RBV1** | **RBV2** | **RBV3** | **RBV4** |
| --- | --- | --- | --- | --- |
| **Width ($\mu\text{m}$)** | 960 | 670 | 480 | 380 |
| **Height ($\mu\text{m}$)** | 1000 | 750 | 650 | 550 |
| **RBV Pressure $P_i$ (Pa)** | -194 | -271 | -358 | -441 |
| **$P_j$ during drainage (Pa)** | -238 | -309 | -390 | -443 |

To experimentally validate our design, we 3D-printed the CC with our calculated dimensions for the main resistor and made PDMS replicas as described earlier (see Fig. 3). Then we tested liquid delivery using aqueous food dye solutions. As expected, each side branch drained sequentially without drainage of the other RBVs (Fig. 4c and Movie S1). Pre-programmed drainage of the side branches was completed within 4 min. The sequence of RBV drainage was 100 % successful over four repeated tests with devices made from four different 3D-printed molds.

#### Capillaric circuit for autonomous delivery of eight liquids

After establishing general guidelines for designing 3D-printed RBVs to obtain sequential liquid delivery, we designed a CC with eight liquid delivery steps, double the number in our proof-of-principle 3D-printed CC and exceeding the number of sequentially-encoded, self-regulated microfluidic drainage events in our previous work with cleanroom-fabricated CCs.^13^

As described earlier, the smallest microchannel width that we could print without a high incidence of defects was 200 μm. We designed the capillary pump region of the 3D-printed CC to be 300 μm wide and 50 μm deep to ensure reliable printing since the capillary pump has a larger pressure than all the RBVs in the circuit. To encode capillary pressure differences, we systematically varied the heights and widths of microchannels in each side branch (see Table 2). We designed RBVs according to the junction pressure criterion (see equation 4) to ensure that valves were drained sequentially. Although in theory, very small differences in capillary pressure between successive RBVs should ensure serial drainage, empirical tests yield that designed capillary pressure differences of ~ 40 Pa provided reliable sequential drainage of RBVs. This empirical value depends on the resolution and accuracy of features produced by the 3D printer and might be reduced with a more accurate printer, or conversely might need to be increased for experiments that require solutions with different surface tensions that will affect the contact angle and the capillary pressure *Pi* of the RBV and branch loaded with this solution.

The main resistor was 18.5 mm long, 300 μm wide, and 50 μm deep to obtain a calculated drainage time of ~10 min for all 8 liquid delivery steps based on the capillary pressures, resistances, and volumes of the microchannels in the circuit. The smallest RBV in the circuit was 380 μm wide and 200 μm deep. Since this valve was much shallower than the reservoir (960 μm wide and 1000 μm deep), we connected the valve to the reservoir using a gently sloped staircase with 50 μm height increments to prevent from liquid stopping due to the formation of an undesired stop valve. We set the maximum channel height in our CCs to 1 mm to stay within a regime where capillary forces are dominant. These geometric constraints limited the number of RBVs, and by extension the number of liquid delivery steps that we could automate in our 3D-printed CCs.

**Table 2.** RBVs designed for CC for autonomous delivery of eight liquids.

|  | **RBV1** | **RBV2** | **RBV3** | **RBV4** | **RBV5** | **RBV6** | **RBV7** | **RBV8** |
| --- | --- | --- | --- | --- | --- | --- | --- | --- |
| **Width ($\mu\text{m}$)** | 770 | 670 | 580 | 480 | 380 | 380 | 380 | 380 |
| **Height ($\mu\text{m}$)** | 900 | 750 | 650 | 600 | 600 | 400 | 300 | 200 |
| **Valve Pressure $P_i$ (Pa)** | -233 | -270 | -314 | -365 | -430 | -483 | -536 | -642 |
| **$P_j$ during drainage (Pa)** | -257 | -295 | -339 | -390 | -455 | -508 | -561 | -645 |

We experimentally validated the operation of the 8-step circuit by 3D-printing a mold and making PDMS replicas as described previously. Fig. 5 shows time-lapse images of autonomous and sequential delivery of 8 liquids in the 3D-printed CC. The autonomous sequential liquid delivery is shown in Supplementary movie S2. Liquids were initially pre-loaded into the reagent reservoirs, and then the central release channel was filled with 10 μL of liquid to start the autonomous drainage operations. Following drainage of the solution from inlet 8 and pinning of the air-liquid interface at RBV8 in the trigger channel, RBV1 is the first to start bursting at t = 3 min 11 s. Each RBV with its attendant reservoir take ~ 50 s to drain and the pre-programmed drainage of the 8 solutions was completed in < 7 min.

**Figure 5.**
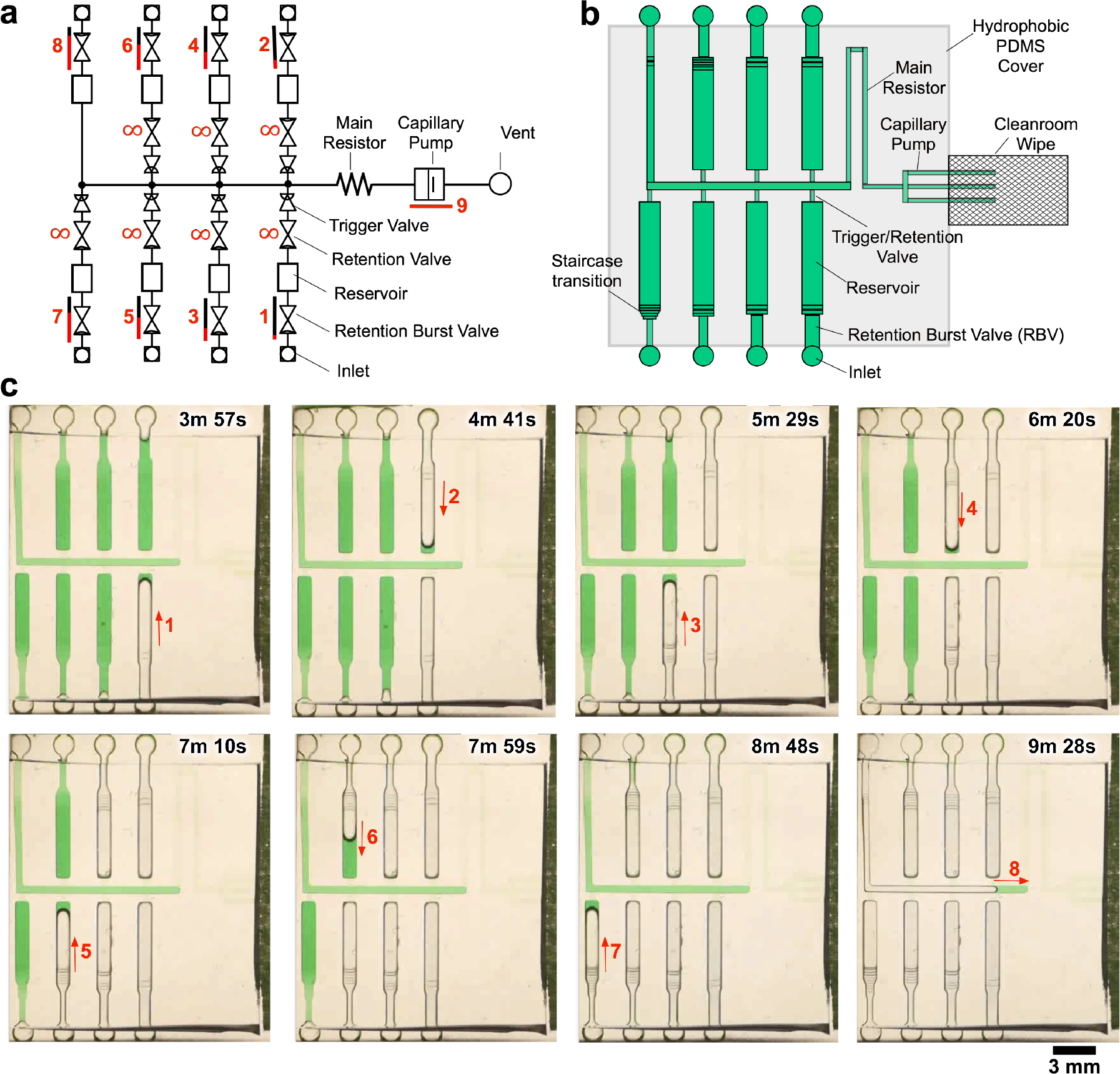
3D-printed CC for autonomous delivery of eight liquids. a) Symbolic representation of CC showing modular assembly of fluidic elements. b) Schematic representation of CC. c) Time-lapse images showing pre-programmed delivery of 8 liquids in 3D-printed CC. Arrows indicate flow direction and numbers highlight the time and sequence of liquid delivery. A video of liquid delivery is provided in supplementary movie S2.

Capillary forces are dominant over gravitational and inertial forces at the scale of the conduits used for the CCs shown here. However, the 3D-printed conduits extend over several tens of mm in some cases, and if a chip is not horizontal, or in the most extreme case if it is positioned such that the channel is vertical, then a hydrostatic pressure will build up that could disrupt functionality of the CC and notably the pre-programmed drainage order. Indeed, the 40 Pa difference between RBVs used here corresponds to a water column of 4 mm in height. Hence, for the reliable operation of CCs with large conduits and incremental differences in capillary pressure it is important to position the devices horizontally to ensure that liquid is routed according to the pre-programmed sequence.

**Table 3.**
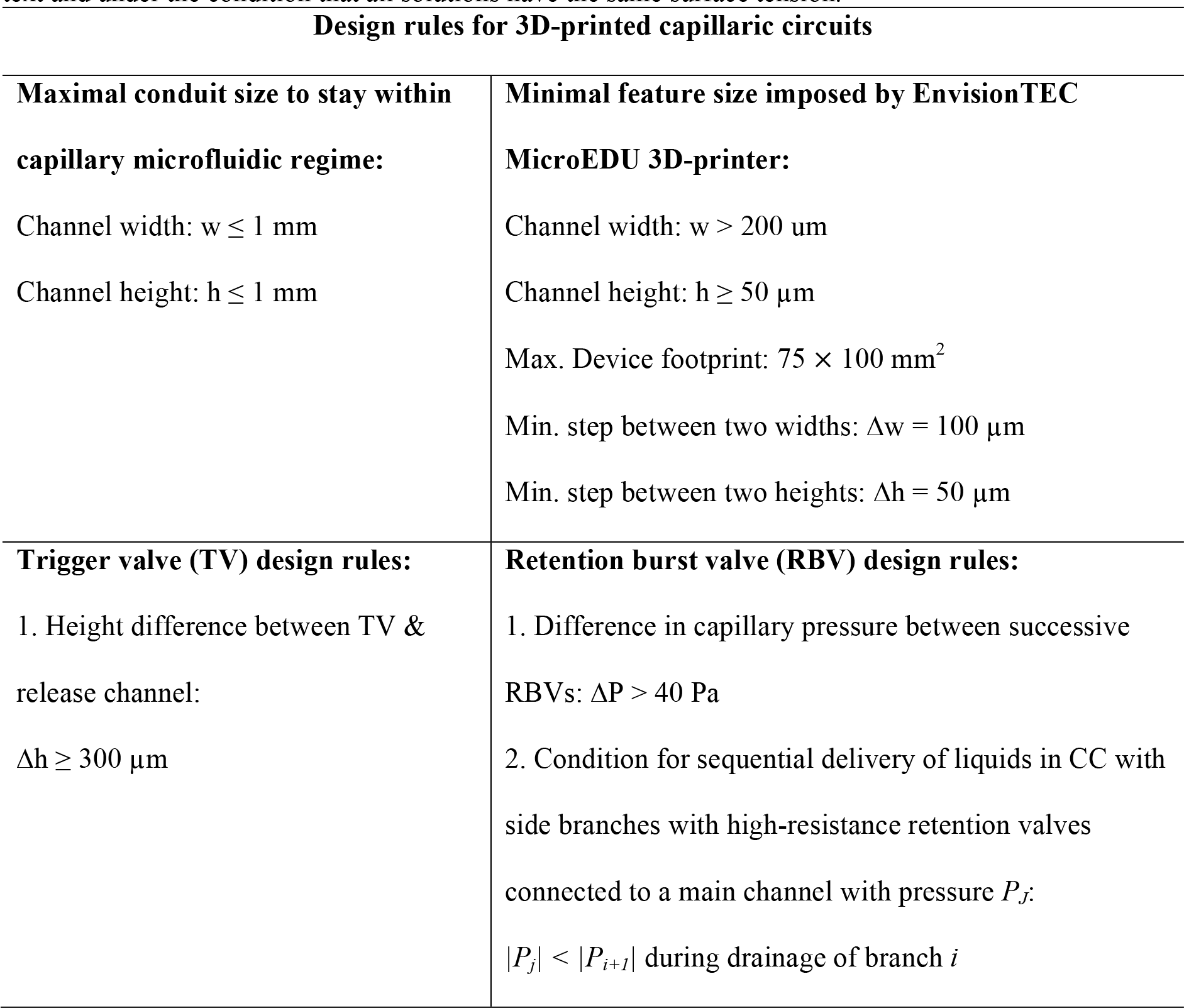
Summary of calculated and empirical design rules for CCs with TVs and RBVs printed using the EnvisionTEC MicroEDU 3D-printer, and replicated into PDMS with surface properties described in the text and under the condition that all solutions have the same surface tension.

## Conclusions

Taken together, our results indicate that 3D-printing allows rapid and inexpensive manufacturing of reliable capillaric valves and circuits with larger volumes and greater functionality than their cleanroom-fabricated counterparts. We characterized the performance of 3D-printed TVs as a function of their geometry. Subsequently, we 3D-printed CCs assembled from fluidic components using an electric analogy. We demonstrated the capabilities of stereolithograpically 3D-printed CCs by autonomously delivering eight liquids in < 7 min, exceeding the number of self-regulated and self-powered liquid delivery steps in our previous work with cleanroom-fabricated CCs by simultaneously changing both width and depth of successive RBVs.^13^ We established design rules for 3D-printed CCs, TVs, and RBVs retention (see Table 3). These design rules are specific to our 3D-printer and the PDMS replicas with a hydrophobic top surface (advancing and receding contact angles of 114° & 89°, respectively) and hydrophilic bottom and side surfaces (advancing and receding contact angles 45° & 31°, respectively). With better 3D-printer accuracy and resolution, more RBVs and liquid delivery steps can be included in the CC; however, the more RBVs there are in the circuit, the more conditions there are to satisfy and the less freedom there is to adjust the resistances (RM, RRV) and pressures (PRV, PC) in the circuit and to adjust flow rates and overall liquid delivery time.

3D-printed CCs are an accessible liquid delivery option that lies between paper microfluidics and cleanroom-fabricated microfluidics. While retaining the deterministic flow control and programmability of cleanroom-fabricated circuits, 3D-printing allows rapid prototyping of new devices within minutes. Currently, the yield of working devices rapidly decreases for features below 200 μm wide. However, as 3D-printers improve, such features and smaller ones may be printed and even more advanced capillary circuits made.

In future, it would be desirable to replace PDMS – which only retains its hydrophilicity for a few hours after plasma treatment^32^ – with alternate polymers with more stable hydrophilic surfaces^33^ and that may be mass produced by injection molding. Alternatively, CCs may be directly 3D-printed into non-transparent or transparent materials with the possibility of using embedded conduits;^34,35^ controlling the surface chemistry is one of the foreseeable challenges that will need to be addressed. With the widespread adoption of 3D-printers, CCs could be readily printed by many researchers, and the design rules presented here will facilitate the fabrication of functional circuits. 3D-printed autonomous CCs may be developed for large-volume and multi-step biochemical assays to be used for point-of-care diagnosis, for research in a lab, as well as for educational purposes.

## Acknowledgements

A.O. acknowledges financial support from NSERC CREATE Integrated Sensor Systems Training Program, CIHR Systems Biology Training Program, and FRQNT PBEEE Scholarship. D.J. acknowledges support from a CHRP grant and from the Canada Research Chair program. We thank the staff at McGill Nanotools Microfab for their microfabrication support. We thank Nicolas Broduch and Professor Raynald Gauvin in the Department of Materials Engineering at McGill University for Scanning Electron Microscopy support. A.O. also acknowledges helpful discussions with Arya Tavakoli, Donald MacNearney, Jeffrey Munzar, and Philippe DeCorwin-Martin at McGill University, as well as Steven Jim at the University of California, Irvine as well. A.O. is grateful for writing feedback from members of his McGill Writing Centre Graphos Peer Writing Group: Ali Moridnejad, Alison Fraser, Krista Oke, Pranavkumar Joshi, Qinglong Wang, and Yevgen Nazarenko.

